# Vestibular and proprioceptive contributions to trunk stabilization differs across postural tasks and walking speeds

**DOI:** 10.1101/2025.06.17.657409

**Authors:** Yiyuan C. Li, Sjoerd M. Bruijn, Koen K. Lemaire, Simon Brumagne, Jaap H. van Dieën

## Abstract

Stabilizing the upright posture of the trunk relies on vestibular and proprioceptive afference. Previous studies found that the feedback responses to sensory afference vary between postures and tasks. We investigated whether and how vestibular and proprioceptive afference contribute to trunk stabilization during different postural tasks, and during walking at different speeds. Twelve healthy adults performed tasks in a random order: sitting, standing on the right foot or both feet, and treadmill walking at five speeds: 0.8, 2.0, 3.2, 4.3 and 5.5 km/h, while exposed to unilateral muscle vibration on the right paraspinal muscles at the level of the second lumbar vertebra, or to a step-like electrical vestibular stimulation (EVS) with the anode behind the left ear. The mediolateral displacements of markers at the sixth thoracic level and sacrum in the global coordinate system were used to evaluate the responses to sensory stimulation. No significant responses to EVS at T6 and sacrum level were found in sitting and standing. Responses to muscle vibration were significant and differed between unipedal standing compared to sitting and bipedal standing. The latter suggests a different interpretation of the sensation of muscle lengthening in these postures. During walking, the magnitude of the responses to both stimuli increased from very slow speeds to moderate speeds. From moderate to higher speeds, responses to muscle vibration decreased, whereas responses to EVS plateaued. These findings suggest speed-dependent modulation of vestibular and proprioceptive contributions in trunk stabilization during walking.

**Summary statement:** By applying electrical vestibular stimulation and muscle vibration, we found that how vestibular and proprioceptive signals are used for trunk stabilization differs between postural tasks and walking speeds.

## Introduction

The trunk represents approximately 50% of the total body mass (de Leva, 1996). With its location high above the base of support, a small deviation from the upright position can produce a substantial destabilizing moment. Trunk muscles have an active role in maintaining the upright posture of the trunk and head (Hof, 2007; Li et al., 2024a; Wu et al., 2019), which supports the use of visual and vestibular information for postural control.

Feedback information for the stabilization of upright posture against gravity is provided by different sensory modalities (Asslander and Peterka, 2014; Maurer et al., 2006; Peterka, 2002; Peterka and Loughlin, 2004). For example, vestibular signals contribute to perception of the head orientation in space (Cullen, 2012), signals from muscle spindles provide a sense of trunk orientation and movement (Prochazka, 2021), and visual information is used to assess the direction and speed of self-motion (Gibson, 1958). These signals need to be integrated to obtain estimates relevant for stabilization of upright posture but are also to some extent redundant. It is believed that redundant sensory signals are integrated based on their relative reliability, in a process called ‘sensory reweighting’ (Asslander and Peterka, 2014; Goar et al., 2025; Oie et al., 2002; Peterka and Loughlin, 2004). This reweighting process allows to compensate for imprecision, loss, or interruption of input from one or more sensory modalities.

The contribution of each sensory modality to stability control is commonly investigated with sensory perturbations (Alberts et al., 2019; Anson et al., 2014; Courtine et al., 2007; Day, 1999; Duclos et al., 2014; Li et al., 2024a). For instance, the contribution of vestibular information has been investigated with electrical vestibular stimulation (EVS) (Alberts et al., 2019; Day, 1999; Li et al., 2024a). The contribution of proprioceptive signals has been investigated with muscle vibration, causing an illusion of muscle lengthening (Goodwin et al., 1972; Inglis et al., 1991; Vizirgianakis et al., 2021).

Previous studies suggest that the gain of feedback responses to sensory afference may depend on the task performed or posture adopted. It is likely that this task- or posture-specific effect of sensory feedback relates to the difficulty of stabilizing these postures (Ali et al., 2003; Britton et al., 1993; Day et al., 1997; Fitzpatrick et al., 1994; Grangeon et al. 2015; Vizirgianakis et al., 2021). For example, in sitting, with a larger base of support, responses to visual, vestibular and proprioceptive perturbations are significantly smaller than in standing (Ali et al., 2003; Grangeon et al., 2015; Vizirgianakis et al., 2021). Additionally, electrical vestibular stimulation induced responses were found to decrease with increased stance width and with the presence of external support (Britton et al., 1993; Day et al., 1997; Fitzpatrick et al., 1994). In bipedal standing, responses to visual perturbations were significantly smaller compared to unipedal standing, a more challenging posture (Hazime et al., 2012). However, Hazime et al. (2012) reported that ankle vibration had less effect in unipedal standing than in bipedal standing. They assumed that the vestibular signal compensates for the decreased use of proprioception through sensory integration (Hazime et al., 2012). This suggests that posture or task-dependent effects may reflect in part reweighting instead of an overall change in feedback gain. From this perspective, the decreased effect of ankle vibration could also be attributed to decreased proprioceptive input from the ankle being compensated by an increased input from the hip and trunk, since corrective actions from the hip and trunk are required to maintain upright posture during unipedal standing (Riemann et al., 2003).

Unlike in standing, which need continuous stabilization, the contribution of sensory signals to stabilization of walking is phasic, as reflected by phase-dependent gains of feedback responses (Blouin et al., 2011; Ivanenko et al., 2000; Li et al., 2024a; O’Connor and Kuo, 2009). The use of sensory signals in stabilization of walking is affected by walking speed; smaller effects of vestibular stimuli were found at higher compared to lower walking speeds (Brandt et al., 1999; Dakin et al., 2013; Fabre-Adinolfi et al., 2018; Jahn et al., 2000; Schniepp et al., 2012). Visual perturbation by means of optic flow was found to have smaller effects in running than in walking, also suggesting a lower contribution of visual signals at higher speeds (Jahn et al., 2001). Based on sensory reweighting, signals from one or more alternative modalities may compensate for the decreased contributions of vestibular and visual signals to stability control during faster walking. Proprioceptive signals could be a potential source of compensation in fast walking, as it has been shown that after removal of proprioceptive feedback, mice maintained the ability to walk at slower speeds but failed to walk at higher speeds (Mayer et al., 2018; Takeoka et al., 2014). However, to the best of our knowledge, similar findings have not yet been reported in humans. Additionally, these studies only included limited speed ranges, i.e. comparing walking at a selected speed to running, or walking at a higher speed.

Here, we studied if, and how, proprioceptive and vestibular signals contribute to trunk stabilization during sitting, unipedal standing, bipedal standing and during walking at different speeds, by using muscle vibration on lumbar paraspinal muscles and electrical vestibular stimulation. We extended the range of walking speeds tested in previous studies to very slow speeds. We focused on the initial responses to step-like sensory stimulation, as later responses will be more affected by the gravitational perturbation that may follow from compensatory movements, and this effect varies between postures. We hypothesized that compared to bipedal standing, during unipedal standing, which poses a greater challenge to stability, both EVS and muscle vibration would lead to larger responses. Similarly, during sitting, which is inherently more robust with a larger base of support, the responses to EVS and muscle vibration would be smaller than during bipedal standing. During walking, EVS was expected to elicit larger responses at slower speeds as reported previously, whereas muscle vibration was expected to elicit larger responses at faster walking speeds because of sensory reweighting.

## Methods

### Participants

Nineteen healthy adults were recruited. Seven participants did not complete all measurements due to a damaged muscle vibrator. In total, data from twelve participants were used in this study (N = 12, 7 males, 5 females, age: 19.9 ± 2.1 years, weight: 65.7 ± 7.5 kg, height: 1.72 ± 0.09 meters). Exclusion criteria for participation were any diagnosed orthopaedic or neurological disorders, or the use of medications that can cause dizziness. All participants provided informed consent. Procedures were approved by the VU Amsterdam Research Ethics Committee (VCWE-2023-136).

### Electrical vestibular stimulation

Electrical vestibular stimulation was applied as an analogue signal through a digital-to-analogue converter (National Instruments Corp., Austin, USA) to an isolated constant-current stimulator (BIOPAC System Inc., Goleta, USA). The current was delivered via two carbon rubber electrodes (9 cm^2^), which were coated with electrode gel (SonoGel, Bad Camberg, Germany) and placed over the mastoid processes with the anode on the left and cathode on the right. In this configuration, a step-like EVS signal, with an amplitude of 1 mA, induces an body sway to the anodal side, i.e. to the left, in bipedal standing (Day, 1999; Lund and Broberg, 1983).

### Unilateral muscle vibration

Two muscle vibrators, consisting of DC motors (Maxon International AG, Sachseln, Switzerland) driving an eccentric mass in a PVC case, were placed 2 cm lateral to the L3 spinous processes on both the left and right paraspinal muscles. Vibrators were fixed tightly with double sided tape and an elastic band. Only the right vibrator was activated (80 Hz, with an amplitude of approximately 2 mm), which induces an illusion of right paraspinal muscle lengthening (Courtine et al., 2007).

### Kinematics and ground reaction force

Whole-body kinematics were recorded using a 3D motion capture system (Optotrak, Northern Digital Inc., Waterloo, Ontario, Canada) sampling at 50 samples/s with cluster markers on the feet, shanks, thighs, pelvis, trunk, head, upper arms and forearms. Corresponding anatomical landmarks were digitized with a six-marker probe based on the model described by Kingma et al. (1996). Ground reaction forces were measured at 1000 samples/s by force plates embedded in the split-belt treadmill (ForceLink b.v., Culemborg, the Netherlands).

### Protocol

For familiarization, participants were exposed to 5-second step-like EVS and muscle vibration during unipedal standing, with three repetitions of each stimulation. Participants then walked on the dual-belt treadmill at the fastest (5.0 km/h) and slowest selected speed (0.8 km/h) for 2 minutes at each speed without any stimulation. The subsequent conditions were defined by the stimulation received (EVS or muscle vibration) and by postural tasks (unipedal standing, bipedal standing, sitting, and walking at five different speeds). The order of conditions was randomized. All conditions were performed with eyes open. With the configurations in this study, EVS typically induces a compensatory body sway to the left, while unilateral muscle vibration typically induces a compensatory body sway to the right during bipedal standing.

During unipedal standing, participants stood on their right foot with arms relaxed at their sides. Trials were repeated if the non-stance leg touched the ground. For bipedal standing, participants stood with their feet shoulder-width apart and arms relaxed at their sides, while for sitting, participants sat upright on a stool with arms crossed at shoulder height, feet on the ground, and knees flexed at 90 degrees. In all postural conditions, participants were instructed to minimize the movement. The 5-second stimulation (muscle vibration or EVS) in standing and sitting trials was manually triggered by the researcher with approximately 10-second intervals before and after stimuli, which aimed to ensure that the participant could not anticipate the stimulus. Each postural task included 10 repetitions of each stimulation. Each repetition lasted 25 seconds in total.

For the walking conditions, participants walked on the dual-belt treadmill at speeds of 0.8, 2.0, 3.2, 4.3 and 5.5 km/h for 4 minutes at each speed. For both EVS and muscle vibration, the stimulation was automatically triggered at right heel strikes and lasted 5 seconds, with approximately 10-second intervals between stimulations.

### Data analysis

Ground reaction forces were down-sampled to 50 Hz. Gait events were identified based on the combined centre of pressure (Roerdink et al., 2008). The mediolateral displacement of markers placed on the T6 spinal process and sacrum were analysed to assess the response to stimulations. A negative mediolateral displacement indicated a leftward movement.

To calculate marker displacement, the trajectories of the T6 and sacrum markers were filtered using a bi-directional 4^th^ order, 10 Hz Butterworth low pass filter. For the walking conditions, the marker trajectories were normalized to the gait cycle starting from right heel strikes. To remove the offset, the baseline, defined as the mean marker position over 2 seconds before the onset of perturbation in standing and sitting, and over one stride before the onset of perturbation in walking, was subtracted from the marker trajectory for each repetition, trial and participant. Marker displacement was then calculated as the mean position during the first 2 seconds after the trigger in standing and sitting tasks, and the first two strides after the trigger in walking. The window size was chosen based on a previous study showing that no turning was induced by EVS during overground walking until the second step, and the initial turn (up to 3 steps) lasted for 2.3 to 3.5 seconds (Fitzpatrick et al., 1999). We also analyzed 1-second and 3□-seconds windows in standing and sitting. It confirmed that the main results were consistent, except for larger effect of perturbations in unipedal standing with the 3-seconds window, which may be due to gravitational effects from compensatory movement. The average displacement was calculated across repetitions for each participant, and then across participants (Fig. 1). Anteroposterior (AP) displacement of the T6 markers were also analyzed. The standard deviations (SD) of T6 displacement in the AP direction were calculated for baselines and perturbed gait cycles at all walking speeds.

**Fig. 1.**
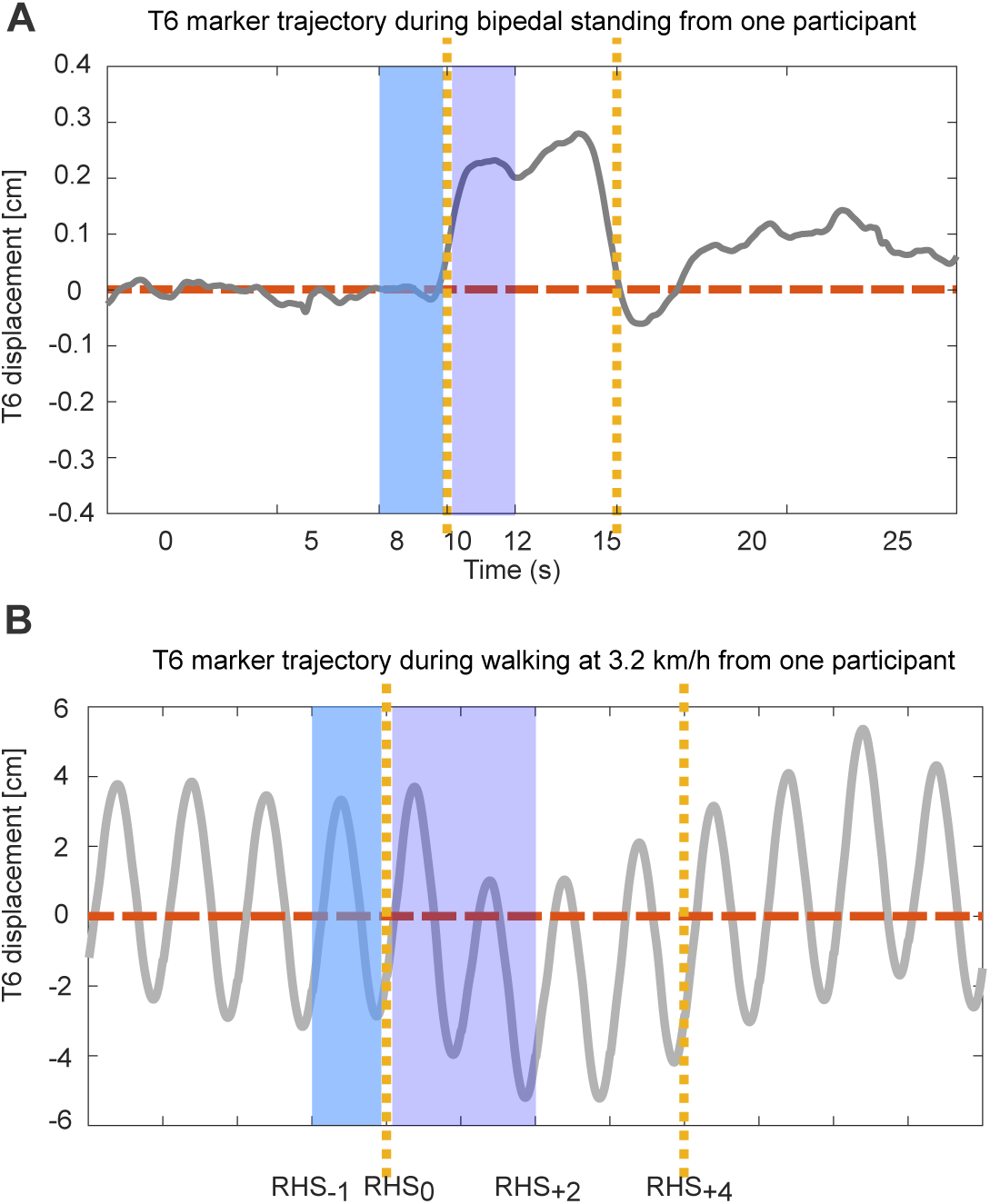
Example of signal analysis. Example of signal analysis of data collected during (A) bipedal standing and (B) walking at 3.2 km/h. Grey lines represent the mean T6 trajectory over ten repetitions for one participant after offset removal. The vertical yellow dashed lines indicate the onset and offset of muscle vibration. Shaded areas represent the baselines (blue) and responses (purple) used for averaging. RHS indicates to right heel strike.

### Statistical analysis

To verify that EVS and muscle vibration predominantly induced perturbations in the mediolateral direction during walking, one-sample t-tests were performed comparing changes in the standard deviation of T6 displacement in the AP direction to zero. To evaluate whether there were effects of EVS and muscle vibration on trunk stabilization, one sample t-tests were performed to compare the mediolateral displacements for each condition to zero. To assess whether the effect of EVS and muscle vibration varied across postures in standing and sitting or speed in walking tasks, we used one-way repeated measured ANOVAs with factors Postural task and Walking speed. Mauchly’s test was used to check the assumption of sphericity, and Greenhouse Geisser corrections were applied when the assumption was rejected. Non-integer values in degree of freedom were due to Greenhouse-Geisser correction. A Bonferroni correction was applied for the post hoc tests when appropriate. A p-value less than 0.05 was defined as significant. Effect size was estimated by Cohen’s d (d) for t-tests and partial eta-squared (η_p_^2^) for ANOVA tests. Statistical analyses were performed in MATLAB (2019a, The MathWorks, Natick, US).

## Results

### Standing and Sitting

EVS induced no significant mediolateral displacement of T6 during unipedal standing (mean ± standard deviation: -0.05 ± 0.24 cm, p = 0.469, d = -0.21), bipedal standing (-0.05 ± 0.15 cm, p = 0.281, d = -0.33), or sitting (-0.01 ± 0.36 cm, p = 0.467, d = - 0.03) (Table 1). There was no significant effect of Postural task on EVS induced displacement of T6 (F (2,22) = 0.242, p_task_ = 0.787, η_p_^2^ = 0.02) (Fig. 2A). Also, no significant mediolateral displacement of the sacrum was induced by EVS during unipedal standing (-0.05 ± 0.16 cm, p = 0.249, d = -0.31), bipedal standing (-0.01 ± 0.14 cm, p = 0.820, d = -0.07), and sitting (-0.00 ± 0.00 cm, p = 0.06, d = 0.00) (Table 1). There was no significant effect of Postural task on EVS induced displacement of the sacrum (F (2,22) = 0.600, p_task_ = 0.588, η_p_^2^ = 0.05) (Fig. 2B).

**Table 1.** Summary of the one sample t-tests on stimulation induced mediolateral displacements during uni-, bi-pedal standing and sitting.

| Postural<br>task | T6 displacement |  |  |  |  |  | Sacrum displacement |  |  |  |  |  |
| --- | --- | --- | --- | --- | --- | --- | --- | --- | --- | --- | --- | --- |
|  | EVS |  |  | MV |  |  | EVS |  |  | MV |  |  |
| | Mean ±<br>SD (cm) | p | d | Mean ±<br>SD (cm) | p | d | Mean ±<br>SD<br>(cm) | p | d | Mean ±<br>SD (cm) | p | d |
| Unipedal | -0.05 ±<br>0.24 | 0.469 | -0.21 | -0.15 ±<br>0.25 | 0.056 | -0.60 | -0.05 ±<br>0.16 | 0.249 | -0.31 | -0.20 ±<br>0.26 | 0.023 | -<br>0.77 |
| Bipedal | -0.05 ±<br>0.15 | 0.281 | -0.33 | 0.10 ±<br>0.11 | 0.008 | 0.91 | -0.01 ±<br>0.14 | 0.820 | -0.07 | 0.03 ±<br>0.07 | 0.136 | 0.43 |
| Sitting | -0.05 ±<br>0.15 | 0.467 | -0.03 | 0.16 ±<br>0.14 | <0.001 | 1.14 | -0.00 ±<br>0.00 | 0.061 | 0.00 | 0.00 ±<br>0.01 | 0.069 | 0.00 |

**Fig. 2.**
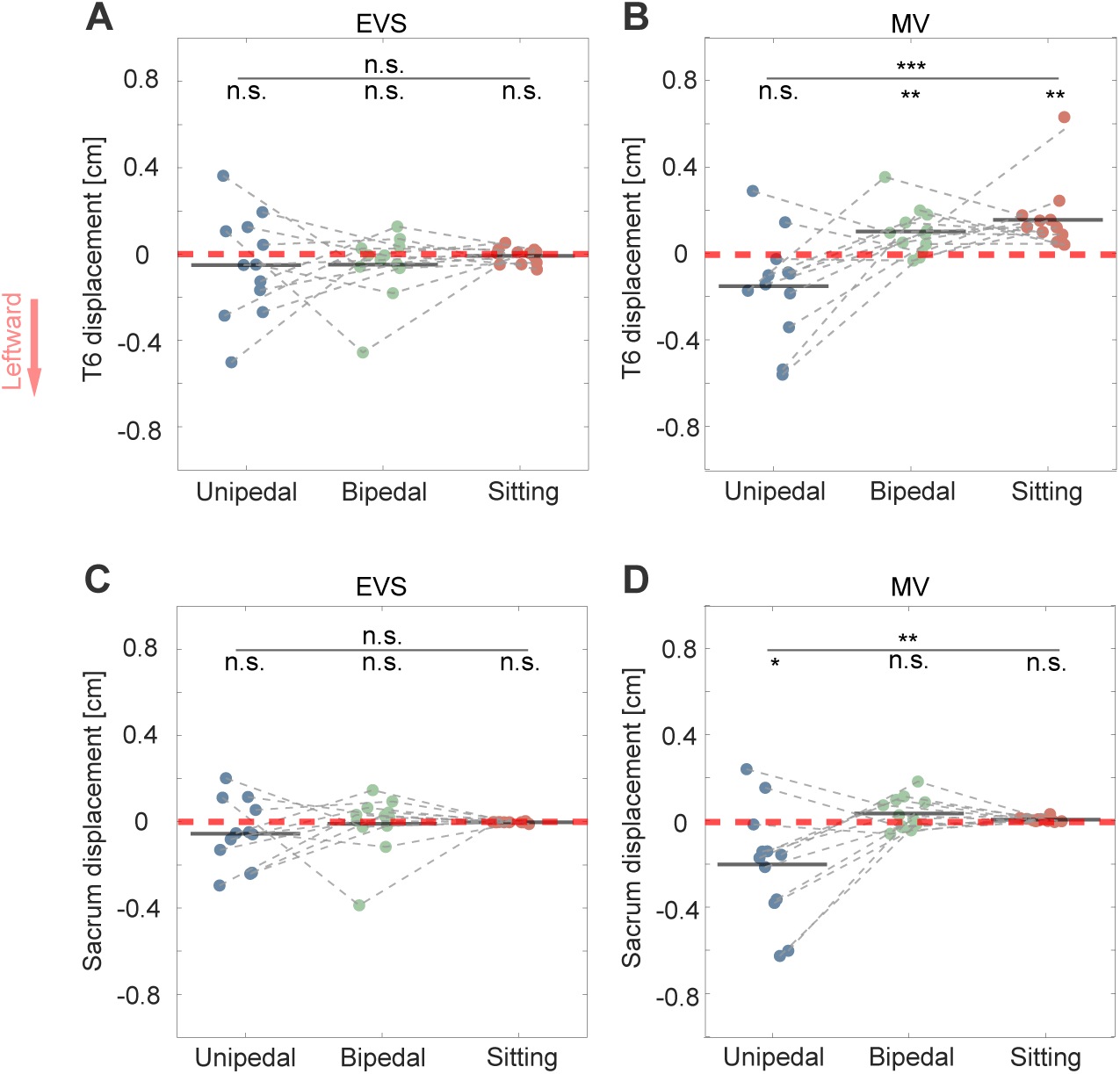
Response during standing and sitting. Mediolateral displacement of T6 (A & B) and sacrum (C & D) in response to electrical vestibular stimulation (EVS, A & C) and muscle vibration (MV, B & D) during unipedal standing (blue dots), bipedal standing (green dots) and sitting (red dots) (N= 13 for all conditions). Negative values (in centimetres, cm) indicate leftward displacements. Group level displacements are shown as black horizontal bars, and individual displacements are represented by filled dots. Muscle vibration induced significant displacement of T6 during bipedal standing (p = 0.008) and sitting (p = 0.003), and significant displacement of sacrum during unipedal standing (p = 0.023; one sample t-test against zero). No significant responses were induced by EVS in any condition. Repeated measured ANOVAs showed that Postural task had no significant effect on EVS induced displacement but significantly affected MV induced displacement of T6 (F (1.686,18.546) = 10.655, p < 0.001) and sacrum (F (2,22) = 8.063, p = 0.002). Asterisks indicate significant p-value, and n.s. indicates non-significance.’

Muscle vibration induced a non-significant leftward displacement of T6 during unipedal standing (-0.15 ± 0.25 cm, p = 0.056, d = -0.60) and a significant rightward displacement of T6 in bipedal standing (0.10 ± 0.11 cm, p = 0.008, d = 0.91) and sitting (0.16 ± 0.14 cm, p < 0.001, d = 1.14) (Table 1). The repeated measured ANOVA test confirmed a significant effect of Postural task on the muscle vibration induced displacement of T6 (F (1.686,18.546) = 10.655, p < 0.001, η_p_^2^ = 0.49,) (Fig. 2A). For the sacrum, muscle vibration induced a significant leftward displacement in unipedal standing (-0.20 ± 0.26 cm, p = 0.023, d = -0.77), but no significant displacement in bipedal standing (0.03 ± 0.07 cm, p = 0.136, d = 0.43), or sitting (0.00 ± 0.01 cm, p = 0.069, d = 0.00) (Table 1). There was a significant effect of Postural task on the muscle vibration induced displacement of the sacrum (F (2,22) = 8.063, p = 0.002, η_p_^2^ = 0.42) (Fig. 2B).

### Walking

No significant effect of either EVS or muscle vibration (MV) were found on the T6 displacement in the AP direction during walking at all speeds, confirming that perturbations were induced predominantly in the mediolateral direction during walking. (p_0.8-evs_= 0.835, d = -0.06; p_0.8-mv_= 0.144, d = 0.45; p_2.0-evs_= 0.189, d = -0.40; p_2.0-mv_= 0.803, d = 0.07; p_3.2-evs_= 0.083, d = -0.55; p_3.2-mv_= 0.608, d = -0.15; p_4.3-evs_= 0.527, d = -0.19; p_4.3-mv_= 0.706, d = -0.11; p_5.5-evs_= 0.109, d = -0.50; p_5.5-mv_= 0.787), d = - 0.08).

EVS and MV induced significant leftward T6 and sacrum displacements at all walking speeds (Table 2). Speed significantly influenced the displacement at both levels, in both stimulation conditions (EVS-T6: F (3.052, 33.572) = 5.707, p_speed_ < 0.001, η_p_^2^ = 0.34; EVS-Sacrum: F (2.976, 32.736) = 4.551, p_speed_ = 0.004, η_p_^2^ = 0.29; MV-T6: F (3.068, 33.748) = 6.304, p_speed_ = 0.002, η_p_^2^ = 0.36; MV-Sacrum: F (3.412, 37.532) = 4.870, p_speed_ = 0.002, η_p_^2^ = 0.41).

**Table 2.** Summary of the one sample t-tests on stimulation induced mediolateral displacements during walking.

| Speed<br>(km/h) | T6 displacement |  |  |  |  |  | Sacrum displacement |  |  |  |  |  |
| --- | --- | --- | --- | --- | --- | --- | --- | --- | --- | --- | --- | --- |
|  | EVS |  |  | MV |  |  | EVS |  |  | MV |  |  |
| | Mean ±<br>SD (cm) | p | d | Mean ±<br>SD<br>(cm) | p | d | Mean ±<br>SD<br>(cm) | p | d | Mean ±<br>SD (cm) | p | d |
| 0.8 | -0.41 ±<br>0.49 | 0.016 | -0.84 | -0.41 ±<br>0.56 | 0.030 | -0.73 | -0.28 ±<br>0.38 | 0.024 | -0.74 | -0.37 ±<br>0.49 | 0.023 | -0.76 |
| 2.0 | -1.56 ±<br>1.37 | 0.002 | -1.14 | -0.89 ±<br>0.46 | <0.001 | -1.93 | -1.07 ±<br>1.09 | 0.006 | -0.98 | -0.72 ±<br>0.39 | <0.001 | -1.85 |
| 3.2 | -1.16 ±<br>1.30 | 0.010 | -0.89 | -1.61 ±<br>0.56 | <0.001 | -2.88 | -0.92 ±<br>1.12 | 0.015 | -0.82 | -1.00 ±<br>0.63 | <0.001 | -1.59 |
| 4.3 | -1.38 ±<br>0.90 | <0.001 | -1.53 | -0.98 ±<br>0.57 | <0.001 | -1.72 | -1.12 ±<br>0.81 | <0.001 | -1.38 | -0.84 ±<br>0.57 | <0.001 | -1.47 |
| 5.5 | -1.09 ±<br>0.92 | 0.001 | -1.18 | -0.44 ±<br>0.53 | 0.016 | -0.83 | -0.77 ±<br>0.71 | 0.003 | -1.08 | -0.34 ±<br>0.55 | 0.050 | -0.62 |

The speed-dependent responses to stimulations (EVS and MV) at both T6 and sacrum levels appeared to follow a non-monotonic pattern. Specifically, the induced mediolateral displacement increased from slow speeds (e.g. from 0.8 to 2.0 km/h), peaked at a moderate speed (around 3.2 km/h), and then decreased at higher speeds (approximately from 4.3 to 5.5 km/h) (Fig. 3). To confirm the non-linearity, post-hoc tests comparing the slowest (0.8 km/h) and fastest speed (5.5 km/h) to the moderate speed (3.2 km/h) were performed with paired t-tests.

**Fig. 3.**
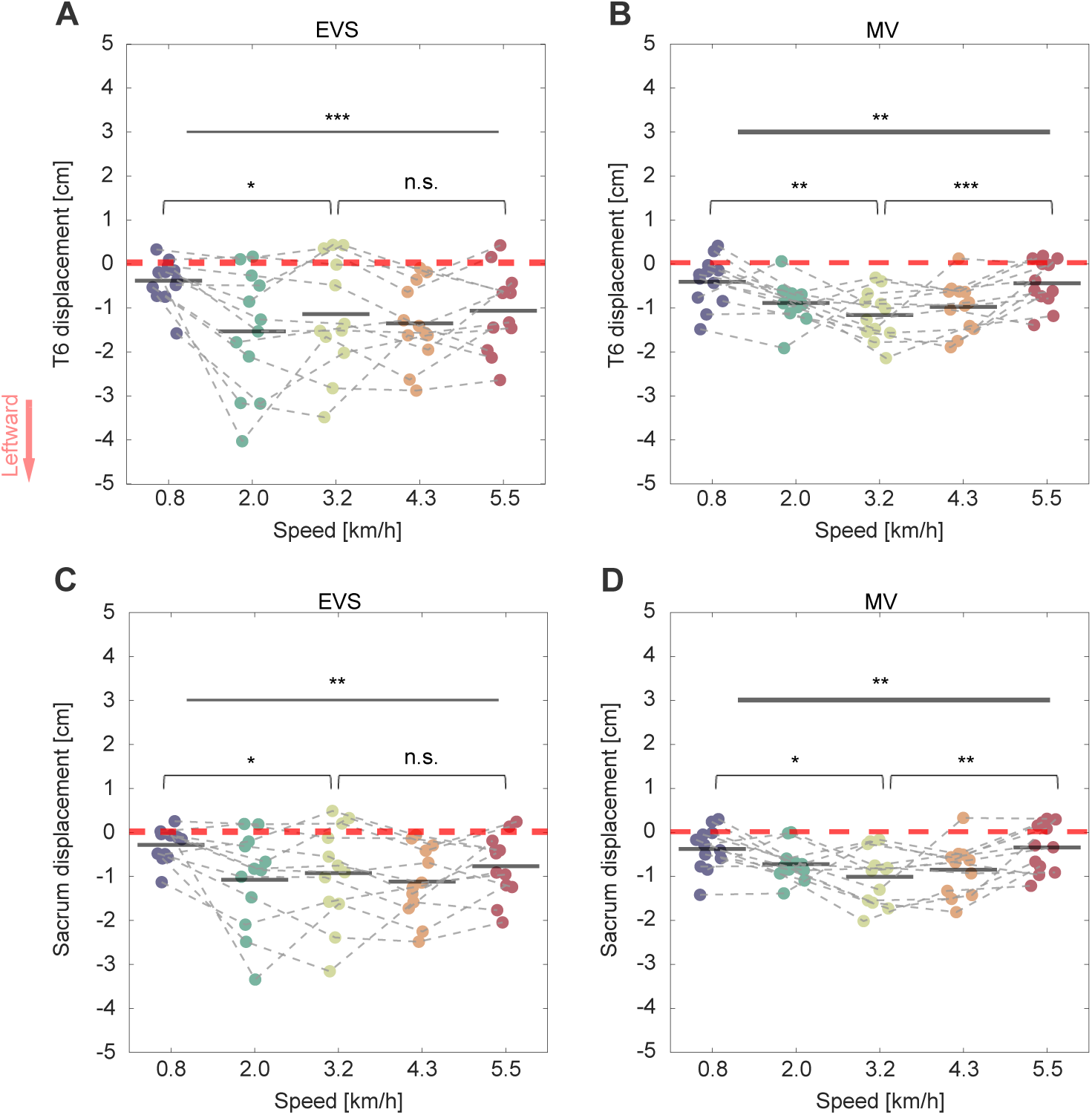
Responses during walking. Mediolateral displacement of T6 (A & B) and sacrum (C & D) in response to electrical vestibular stimulation (EVS, A & C) and muscle vibration (MV, B & D) during walking at five different speeds (N= 13 for all conditions). Negative values (in centimetres, cm) indicate leftward displacements. Group level displacements are shown as black horizontal bars, and individual displacements are represented by filled dots. Significant effects of Speed were found on displacement in response to EVS at both T6 and sacrum level (repeated measured ANOVAs). In the MV conditions, induced responses also varied significantly with speed at T6 and sacrum. Post hoc tests indicated that responses at the slowest (0.8 km/h) speed were significantly smaller than those at the moderate speed (3.2 km/h) in all conditions (paired t-test). Responses at the highest speed (5.5 km/h) were also significantly smaller than at than those at the moderate speed (3.2 km/h) in the MV conditions, but not in the EVS conditions. Asterisks indicate significant p-value, and n.s. indicates non-significance.

Responses to muscle vibration at both T6 and sacrum significantly increased from the slowest to moderate speed (T6: p = 0.004, d = 1.04, sacrum: p = 0.017, d = 0.81) and then decreased at the highest speed (T6: p < 0.001, d = –1.31; sacrum: p = 0.005, d = –0.99), consistent with a U-shaped pattern.

Responses to EVS at T6 and sacrum significantly increased from the slowest to the moderate speed (T6: p = 0.032, d = 0.71; sacrum: p = 0.045, d = 0.65), whereas no significant differences were found between the moderate and fasted speed (T6: p = 0.704, d = –0.11; sacrum: p = 0.402, d = –0.25). Therefore, EVS induced responses increased from very slow to moderate walking speeds and plateaued at higher speeds.

## Discussion

We investigated the task specific contribution of the vestibular and lumbar proprioceptive systems to trunk stabilization during sitting, unipedal and bipedal standing, and walking at five different speeds. Contrary to our hypothesis, EVS did not evoke significant mediolateral displacements during sitting, bipedal standing, or unipedal standing. Muscle vibration evoked task-dependent responses, i.e. it induced a significant leftward displacement of the sacrum during unipedal standing, whereas evoking a significant rightward displacement at T6 only during bipedal standing and sitting. During walking, both EVS and muscle vibration consistently induced significant leftward displacements of T6 and sacrum across all speeds. In line with our hypothesis, the magnitude of these responses varied with walking speed. Below, we discuss how these findings advance our understanding of the task specific contribution of the vestibular and lumbar proprioceptive systems to the trunk stabilization.

### EVS induced responses during standing and sitting

No significant responses to EVS at T6 and sacrum levels were found during sitting, uni- and bipedal standing. This contrasts with previous studies, which observed a significant displacement of centre of pressure, head, torso and pelvis towards the anodal side in response to EVS during bipedal standing (Day et al., 1997; Fitzpatrick et al., 1999; Lund and Broberg, 1983). Our null effect could be explained by the availability of visual information. Previous studies were performed with eyes closed. Hence, in such a condition, vestibular input was likely upweighted, increasing the effect of EVS. Consistent with this, EVS-induced motor responses during bipedal standing were smaller with eyes open compared to closed (Britton et al., 1993; Fitzpatrick and Day, 2004). Furthermore, a study showed that during bipedal standing on a stable surface, reliance on ankle proprioception was greater than on vestibular signals, and this reliance increased with the presence of EVS (Cenciarini and Peterka, 2006). Therefore, even though our sample size and stimulation intensity were comparable to previous studies with eyes closed (Day et al., 1997; Fitzpatrick et al., 1999; Lund and Broberg, 1983), the intensity of EVS may not have been sufficient to evoke a significant effect on trunk stabilization during uni- and bipedal standing due to sensory reweighting.

However, another study reported that the presence of vision did not significantly modulate the EVS induced lumbar muscle responses during sitting (Amélie et al., 2021). The authors attributed this to the strong tactile input from the thighs and buttocks during sitting. Reweighting between vestibular and tactile information has been confirmed by a reduced coupling between EVS and the centre of pressure trajectory during bipedal standing when providing fingertip contact (Goar et al., 2025). From a biomechanical point of view, the large contact of the thighs and pelvis with the seat provides a larger base of support compared to standing. The height of the centre of mass relative to the base of support is lower during sitting, consequently a smaller destabilizing torque will be generated when the trunk has a given angular deviation from the upright posture compared to standing. For these reasons, the gain of responses to EVS induced illusions is likely to be smaller during sitting than during uni- and bipedal standing.

### EVS induced responses during walking

During walking, we found that at both T6 and sacrum level, EVS induced a significant displacement to the left, i.e. the anodal side, in line with previous literature (Bent et al., 2004; Fitzpatrick et al., 1999; Jahn et al., 2000; Iles et al., 2007). The vestibular signal encodes the head orientation in space (Cullen, 2012), and this perceived vestibular signal needs to be integrated with neck proprioception to distinguish whether the whole body is moving or only the head is moving on the stationary trunk (Pettorossi and Schieppati, 2014). In our case, participants were walking facing forward, thus head orientation was on average aligned with the trunk. Consequently, EVS induced an illusion of whole body movement to the right. A global compensatory response to the left was induced to stabilize the trunk in the upright orientation in space.

The differences in the effects of EVS between standing and walking were partly supported by Fitzpatrick et al. (1999), who reported that EVS induced body sway during standing was smaller and more variable than during walking. During walking, a stereotyped and symmetric locomotion, the contribution of vestibular signals is phasic (Blouin et al., 2011; Ivanenko et al., 2000; Li et al., 2024b; O’Connor and Kuo, 2009). As such, the phase-locked step-like EVS induced perturbation is likely to be consistent over strides. This phasic vestibular contribution was also reported in animals, with enhanced responses to vestibular stimulation at the initial stance phase, presumably to provide antigravity support and maintain the equilibrium of the trunk (Marlinsky, 1992). Biomechanically, vestibular signals are selectively used during phases where stabilization demands are greatest due to changes in base of support (Fitzpatrick et al., 1999; Magnani et al., 2021). In contrast, during standing and sitting the responses to EVS can superimpose negatively or positively on the ongoing body sway, which would increase the variability in measured responses. In addition, the larger and more consistent responses also suggest increased weighting of vestibular signal, likely reflecting its role in controlling heading direction during walking. It has been shown that, compared to quiet standing, EVS induced responses increased when subjects voluntarily tilted their bodies laterally or forward while standing (Cauquil and Day, 1998; Smetanin et al., 1988). This increase was attributed to the contribution of vestibular signals to spatial perception during voluntary movements (Cauquil and Day, 1998).

### Muscle vibration induced responses during standing and sitting

Muscle vibration causes an illusion of lengthening of the vibrated muscle (Goodwin et al., 1972). Consistent with literature, during bipedal standing and sitting, a significant rightward T6 displacement was found (Courtine et al., 2007). No significant sacrum displacement was found in these tasks. In contrast, during unipedal standing a sacrum displacement, but no trunk displacement, was observed. We therefore analysed the relative position of T6 to sacrum (Fig. 4A). A rightward displacement of T6 relative to the sacrum during unipedal standing was found in all three postures. Thus, the different response during unipedal standing versus sitting and bipedal standing could be caused by a difference in the interpretation of the postural change related to trunk muscle lengthening. During bipedal standing and sitting, the sacrum is likely to maintain stable, given the wide and symmetrical support. Thus, muscle lengthening is likely to be interpreted as a leftward trunk movement relative to the sacrum, inducing a compensatory rightward trunk movement. However, in unipedal standing with a less stable and asymmetric support of the pelvis, this perceived lengthening is possibly to be interpreted as a rightward displacement of the pelvis, inducing a leftward sacrum displacement to re-align the segments.

**Fig 4.**
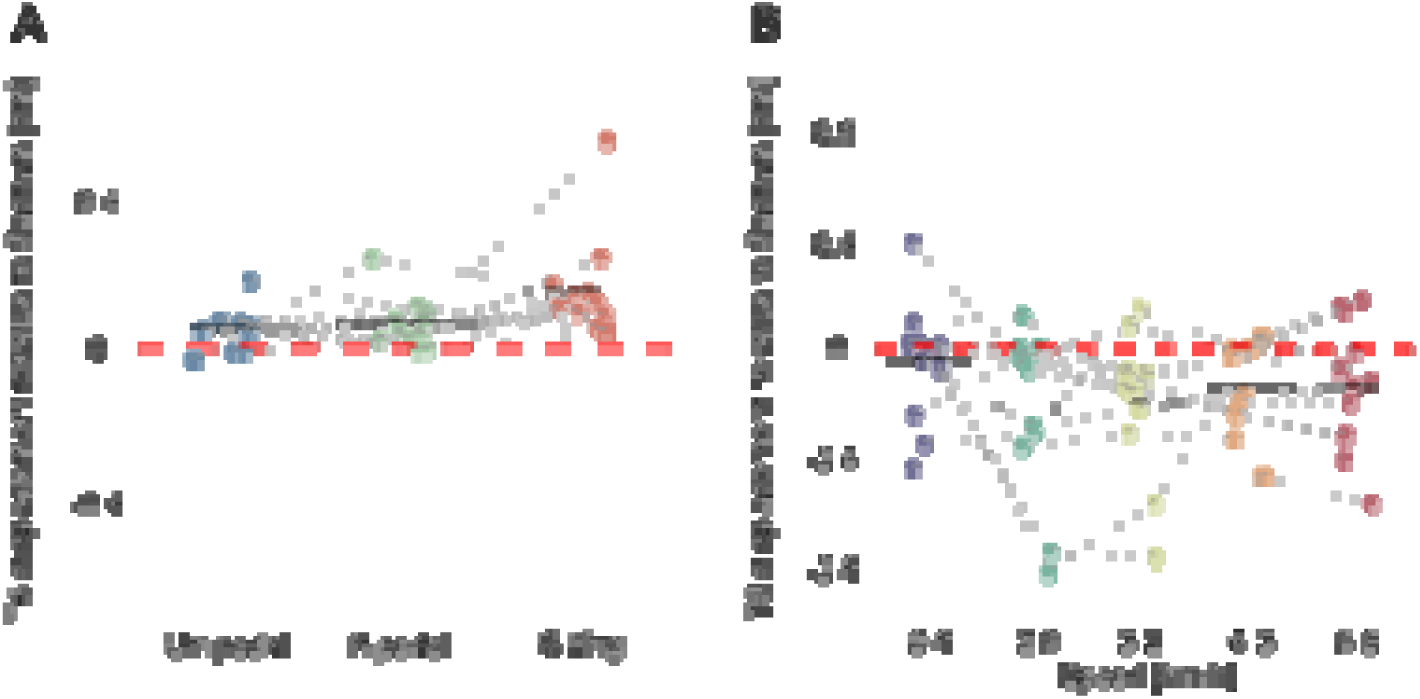
MV induced T6-sacrum relative displacement. MV induced relative displacement between T6 and sacrum in the mediolateral direction across tasks: A) during unipedal standing (blue dots), bipedal standing (green dots) and sitting (red dots); and B) during walking at five difference speeds. Negative values indicate leftward displacement of T6 relative to sacrum (in centimetres, cm). Group level displacements are shown as black horizontal bars, and individual displacements are represented by filled dots.

### Muscle vibration induced responses during walking

Similar to unipedal-standing, during walking, the vibration induced perception of paraspinal muscle lengthening may be interpreted as a pelvis movement. With vibration on the hip abductor muscle of the stance leg, an inward placement of the contralateral foot was reported, which suggests that this vibration similarly induced an illusion of pelvis displacement towards the stance side (Arvin et al., 2018). In line with this, we found that lumbar muscle vibration induced a significant leftward displacement at T6 and sacrum levels opposite to the vibrated side, also consistent with previous studies using trunk muscle vibration (Courtine et al., 2007; Schmid et al., 2005). Unlike during unipedal standing, the T6 displacement was more to the left relative to the sacrum during walking (Fig. 5B). Thus, these responses during walking seem not to be the result of balance control, for which we expected a rightward displacement of T6 relative to the sacrum as found in standing and sitting.

Unilateral lengthening of paraspinal muscles can result not only from trunk or pelvis displacement in the frontal plane, but also from a change in orientation in the transverse plane. Schmid et al. (2005) found that during overground walking with either eyes opened or closed, lumbar paraspinal muscle vibration evoked a deviation in walking trajectory. Additionally, deviations in walking direction were evoked by vibration applied to abdominal muscles and paraspinal muscles at thoracic and lumbar levels, but not with vibrations on lower limb muscles, such as hip abductors and adductors (Courtine et al., 2007). Therefore, trunk proprioceptive information appears to play a major role in the control of balance as well as the control the heading direction during walking.

Interestingly, muscle vibration induced local responses in static conditions but global responses during walking. This likely reflects that proprioceptive inputs from multiple sources are integrated, while the interpretation in terms of an induced illusion depends on the functional role of proprioception in different tasks and postures. This is consistent with a previous study showing that bilateral neck muscle vibration evoked local head movement when trunk was supported but induced whole-body sway without trunk support during sitting and standing (Smetanin et al., 1993).

### Effects of walking speed

The effect of EVS and muscle vibration on trunk stabilization varied with walking speed. Contrary to our hypotheses, in the selected range of speeds from 0.8 to 5.5 km/h, responses to both stimuli increased from very slow speeds to moderate speeds. This may be in line with the phenomenon that specific patient groups with sensory loss or dysfunction tend to walk slower (Hicks et al., 2017; Menz et al., 2004; Wuehr et al., 2014). Some studies suggested that slow walking is more stable than fast walking (Dingwell and Marin, 2006; England and Granata, 2007). However, this conclusion has been debated (Best and Wu, 2020; Bruijn et al., 2009; Hak et al., 2012).

Smaller feedback responses at very low speeds may be a result of the increased duration of the stance phases, especially of the double support phases, both in absolute sense and as a percentage of the gait cycle (Wu et al., 2019; Wuehr et al., 2014). Given the long duration of the double support phases in very slow walking (e.g. 0.8 km/h), the control strategy may resemble that during standing, where we found smaller or no effects of sensory stimulation. However, previously we found that trunk muscle modulation in response to vestibular afference mainly took place during the double support phase (Li et al., 2024a).

Another possible explanation for the smaller feedback responses at very low speeds is that walking at such slow speed is controlled as a repeated transition between standing and walking. The optimal feedback control theory suggests that to enable the transition from one posture or task to another, the control set for the preceding needs to be disengaged before setting sensory feedback gains for the subsequent task (Cluff and Scott, 2016). This prediction was supported by findings from Tisserand *et al*. (2018),who reported absence of coupling between EVS and ground reaction force during the transition between standing and walking. Hence, when very slow walking is regarded as a repeated transition between standing and walking, the overall feedback gains should be low.

From moderate to higher walking speeds, responses to EVS plateaued rather than decreased, in contrast to previous reports of reduced EVS effects at higher locomotion speeds (Brandt et al., 1999; Dakin et al., 2013; Fabre-Adinolfi et al., 2018; Jahn et al., 2000). This discrepancy may be partly explained by differences in experimental design, as most of these studies compared walking with running (Brandt et al., 1999; Fabre-Adinolfi et al., 2018; Jahn et al., 2000). Biomechanically, running differs from fast walking by the absence of a double support phase, during which vestibular afference mostly contributes to trunk muscle modulation during walking (Li et al., 2024a). Therefore, reduced EVS effects observed during running than walking may reflect changes in gait mechanics across the transition between walking and running rather than a monotonic effect in vestibular contribution with increasing speeds in walking.

Dakin et al. (2013) found reduced EVS evoked responses in lower limb muscles when walking at 0.8 m/s (2.9 km/h) compared to 0.4 m/s (1.4 km/h). These speeds fall within the interval between speed conditions in the current study (0.8, 2.0 and 3.2 km/h). Moreover, they applied stochastic GVS signals rather than the step-like EVS. These methodological differences may also contribute to difference finding between current and previous studies. In addition, our statistical power could be limited as there was a small effect size and we had a limited sample size. Therefore, a slight reduction in responses to EVS at higher speed cannot be excluded based on the current data.

Based on previous studies of reduced effects of vestibular and visual perturbations on walking stabilization at higher walking speed (Brandt et al., 1999; Jahn et al., 2001; Dakin et al., 2013; Fabre-Adinolfi et al., 2018; Jahn et al., 2000), we expected that the proprioceptive signal would compensate during fast walking, reflected by a larger muscle vibration induced response was expected with increased walking speed. However, response to muscle vibration decreased from moderate to higher speeds. This finding is consistent with the reduced activity of the vestibular and somatosensory cortex during imagined running compared to imagined slow speed walking and standing (Jahn et al., 2004), suggesting a decreased reliance on feedback control at higher speeds. This interpretation can be further supported by a recent study showing less energetic cost of walking stabilization at fast walking compared to a slow walking (Muijres et al., 2026).

It is believed that stable walking is achieved by the interaction of multisensory feedback control, automated central pattern generators, and modulation of intrinsic mechanical properties of the body (Grillner and El Manira, 2020; MacKay-Lyons, 2002; Wuehr et al., 2013). In animal models without movement-related sensory feedback, automated central pattern generators can generate the rhythmic muscle activity for locomotion (Dubuc et al., 1986; Grillner et al., 1976; Lambert et al., 2012). It is therefore plausible that in human at faster walking speeds where locomotion becomes more rhythmic, the automated locomotor control partially replaces feedback control. Additionally, the decreased effects of muscle vibration at higher speeds may also be explained by biomechanical factors. At faster walking speeds, trunk muscle activity is increased (Anders et al., 2007), which would result in increased muscle stiffness and consequently increased trunk stiffness.

### Limitations

In this study, we did not statistically compare the contribution of vestibular and proprioceptive information between the static and dynamic tasks in this study for several reasons. Dependent on the task, sensory information may be used not only for balance control. As we argue above, trunk proprioceptive information also contributes to the control of walking direction. Moreover, the responses to stimulation were analysed differently between walking and static conditions. Additionally, differences in trunk control strategies could further complicate comparisons between postures and tasks. For example, the responses to muscle vibration at the trunk and pelvis level differ between unipedal standing, bipedal standing and sitting. In addition, the demands on trunk control differ between postures. For example, trunk deviations generate smaller destabilizing torques during sitting than during standing because the CoM ls located lower relative to the base of support. Together, these factors could confound the comparison of response magnitudes to stimulation, i.e. the gain of feedback response, between static and dynamic tasks.

Another limitation is that we cannot investigate whether the relative weights of the vestibular and proprioceptive signals differed between tasks and postures for trunk stabilization. We found parallel trends in the task and speed-related modulation of responses evoked by electrical vestibular stimulation and muscle vibration, which possibly indicates an overall modification of the feedback control gain. At the same time, reweighting between vestibular and proprioceptive signals could also exist for optimal feedback control in different task and postures. However, experimentally quantifying sensory weighting is challenging. Although EVS and muscle vibration provide a direct perturbation to the vestibular system and proprioception, the resulting balance responses likely represent integrated multisensory feedback, in which sensory reweighting due to sensory perturbations may already have occurred. Combining experimental approaches with computational modelling provides a powerful tool to quantify changes in sensory weights in unperturbed conditions.

Within neuromusculoskeletal models, each sensory modality can be independently adjusted by changing parameters such as noise amplitudes, and the effects on balance performance can be evaluated with simulations (Wouwe et al., 2022). We performed two separate repeated-measures ANOVAs for the T6 and sacrum markers to evaluate the effects of EVS or muscle vibration across postural tasks. This may increase the risk of Type I errors by testing the dataset from T6 and sacrum independently, as biomechanically, the motions of T6 and sacrum are not independent. Considering the Marker Position as a factor in the MANOVA test could be a reasonable solution. However, if a significant effect of marker position would be found, ANOVAs for each marker should be performed as post-hoc tests. In this case, the main finding should remain the same as current findings, i.e. the contribution of vestibular and proprioceptive information in trunk stabilization differs between postural tasks and walking speeds.

To test our hypotheses, we averaged marker displacements across repetitions to extract the consistent directional responses to muscle vibration and EVS across postures and tasks. However, averaging does eliminate temporal information on the responses. With the current dataset, future work could access how responses change across repetitions to provide insight into potential adaptation and dynamic reweighting of sensory inputs.

### Conclusion

In conclusion, the present findings confirm that the contribution of vestibular and proprioceptive afference to the trunk stabilization varies between postures and walking speeds. Muscle vibration induced different response at the T6 and sacrum level comparing unipedal standing to sitting and bipedal standing. This suggests a different interpretation of the sensation of muscle lengthening in these postures. By extending the range of selected walking speeds compared to previous studies, we found decreased responses to stimulation at very slow speeds. This suggests that a different control strategy is used during slow walking, possibly due to the long double stance duration, or perhaps because very slow walking reflects a repeated initiation of walking rather than a steady-state process. From moderate to higher speeds, response to EVS plateaued, which could be due to methodological differences to literatures and limited effect size in the current study. Therefore, a slight reduction in responses to EVS at higher speed cannot be excluded. However, response to muscle vibration significantly decreased. Overall, these findings suggest that stabilization of faster walking relies less on feedback control.

## Additional information

### Data Availability Statement

The original data are available from the corresponding author upon reasonable request.

### Competing Interests

The authors declare that there are no conflicts of interest.

### Author Contributions

Experiments were performed at the Dual Belt Lab in the Vrije Universiteit Amsterdam. Conception and design: Y.C.L., K.K.L., S.M.B., S.B., and J.H.D. Data acquisition: Y.C.L. Analysis and interpretation: Y.C.L., K.K.L., S.M.B., S.B. and J.H.D. Drafting and revising manuscript: Y.C.L., K.K.L., S.M.B., S.B. and J.H.D. All authors have read and approved the final version of this manuscript and agree to be accountable for all aspects of the work in ensuring that questions related to the accuracy or integrity of any part of the work are appropriately investigated and resolved. All persons designated as authors qualify for authorship, and all those who qualify for authorship are listed.

### Funding

Y.C.L. was funded by a scholarship (No. 202108520034) from the China Scholarship Council (CSC). S.M.B. was funded by a VIDI grant (no. 016.Vidi.178.014) from the Dutch Organization for Scientific Research (NWO).

## Acknowledgements

The authors appreciate all participants for their voluntary participation. The authors would like to thank Yaqi Li and Yang Geng for their help in data collection. The authors thank Bert Clairbois and Richard Casius for technical assistance.

## Notes

### Competing Interest Statement

The authors have declared no competing interest.

### Summary of Updates

All figures were updated for better visualization. Effect sizes were reported using Cohens d for t-tests and partial eta-squared for ANOVA tests. Discussion and conclusions were modified based on updated statistical analysis.

